# Machine Boss: Rapid Prototyping of Bioinformatic Automata

**DOI:** 10.1101/2020.02.13.945071

**Authors:** J. Silvestre-Ryan, Y. Wang, M. Sharma, S. Lin, Y. Shen, S. Dider, I. Holmes

## Abstract

**Motivation:** Many C++ libraries for using Hidden Markov Models in bioinformatics focus on inference tasks, such as likelihood calculation, parameter-fitting, and alignment. However, construction of the state machines can be a laborious task, automation of which would be time-saving and less error-prone.

**Results:** We present Machine Boss, a software tool implementing not just inference and parameter-fitting algorithms, but also a set of operations for manipulating and combining automata. The aim is to make prototyping of bioinformatics HMMs as quick and easy as the construction of regular expressions, with one-line “recipes” for many common applications. We report data from several illustrative examples involving protein-to-DNA alignment, DNA data storage, and nanopore sequence analysis.

**Availability and Implementation:** Machine Boss is released under the BSD-3 open source license and is available from http://machineboss.org/.

**Contact:** Ian Holmes

## 1 INTRODUCTION

Bioinformatics is a field littered with state machines, many of them still functional. The venerable Needleman-Wunsch, Smith-Waterman, and Gotoh algorithms from the 1970s and early 1980s can be thought of as aligning pairs of sequences to input-output automata [35, 44, 16]. Gribskov’s protein profiles of the late 1980s are state machines too [18]. The 1990s saw the probabilistic interpretation of both these kinds of machine as Hidden Markov Models (HMMs); respectively, the “pair HMM” and the “profile HMM” [7, 12]. This inspired new applications of HMMs in emerging areas of sequence analysis, such as computational gene prediction [9]. The 2000s saw further evolution of these ideas including HMMs for multiple sequence alignment [20, 11], comparative genefinding [2, 32], and phylogenetics [40, 41, 45]. HMMs (and automata more generally) continue to represent the state of the art for many bioinformatic tasks; for example, when reconstructing the indel histories of ancestral sequences [30, 49, 21] or aligning protein to DNA [3]. Meanwhile, the growing field of Deep Learning drew from HMM model-fitting algorithms to train Recurrent Neural Networks (RNNs) with sequential inputs and outputs; specifically, using *Connectionist Temporal Classification* (CTC) which is based on the Forward-Backward algorithm [17]. Inevitably, RNNs have found application in bioinformatics, sometimes surpassing HMMs; for example, RNN basecallers for nanopore sequence data [6] have been shown to outperform the corresponding HMMs [10]. Even in cases such as this, where RNNs have displaced HMMs, useful connections to automata theory can sometimes still be made due to the underlying parallels between CTC and HMM dynamic programming [42].

Over this period, a number of software libraries have been developed for parsing, annotating, and aligning biological sequences using generic state machines. Examples include Dynamite [4], DART [20], GHMM [39], C4 [43], HMMoC [31], HMMConverter [27], StochHMM [29], MuxStep [48], and ham [36]. These various libraries all have slightly different capabilities but typical features include the ability to work with state machines of arbitrary topology, implementations of common dynamic programming algorithms (like the Viterbi and Forward-Backward algorithms), and generation of optimized code implementing those algorithms. Some newer libraries for deep learning that are frequently used in bioinformatics, such as TensorFlow [1], often include implementations of algorithms with close state-machine analogs, even though the libraries themselves are not explicitly founded on automata theory. Examples include CTC loss-minimization, or beam search to find the most likely output sequence of a RNN.

Despite all of these libraries for doing automata-based inference on sequences, there are few (if any) general-purpose tools for working with the automata themselves as manipulable mathematical objects. As one illustration of why this is useful, we consider GeneWise, one of the most successful (and elaborate) automata of bioinformatics. The underlying state machine of GeneWise aligns an amino acid sequence to the (unspliced and imperfectly-observed) protein-coding genomic DNA. In doing so, it simultaneously models translation, splicing, and sequencing errors—as detailed in the GeneWise paper [3]. The machine developed to do this in full is highly intricate, containing 21 states and 93 transitions when expressed as a Moore machine [34]; prototyping and subsequent optimization yielded a reduced-size machine with 6 states and 23 transitions. (In using these descriptions to assess the mathematical complexity of GeneWise and the computational complexity of its algorithms, one should note that the transition output labels are strings, not just individual characters, and the transition weights include contributions from all the constituent sub-models: translation, splicing, and sequencing errors.) GeneWise also includes an algorithm that aligns profile HMMs of protein domains to genomic DNA, accepting HMMER-format profiles as used by the PFAM database [14]. All these state machines were laboriously developed, validated and debugged by hand (albeit with the help of Dynamite to generate the dynamic programming code once the state machines were specified). In principle—given that the GeneWise model can notionally be “factorized” into independent component machines for translation, splicing, and sequencing error—this approach should be amenable to incremental variations or enhancements; e.g. different profile HMM architectures (as may be found in later releases of HMMER), alternate genetic codes, or richer models of the context-dependent error profile of later-emerging sequencing technologies like that of Oxford Nanopore Technologies (ONT) [23]. In practice, however, because this factorization of the GeneWise state machine into sub-machines is performed manually, these kinds of upgrade would take a significant amount of work—and quick prototyping would be impossible.

More generally, many bioinformatic automata can be viewed as being derived from simpler state machines by operations such as concatenation, composition (or multiplication), intersection, union (or addition), reversal, complementation, substitution, or other well-defined mathematical transformations. This is particularly useful in cases which inherently involve transforming one sequence into another via several steps. GeneWise is one example. Another example occurs in the context of encoding binary information into DNA as a storage medium: in doing this, it is desirable to avoid repeated nucleotides (which are easily misread by DNA sequencers) and this can be conceived of as converting a binary sequence to a base-3 (ternary) sequence, followed by a conversion from ternary to DNA. Each conversion can be formulated as an input-output state machine (Figure 1); related coding operations, such as the introduction of parity bits for error correction, can similarly be formulated using state machines (Figure 2).

**Fig. 1.**
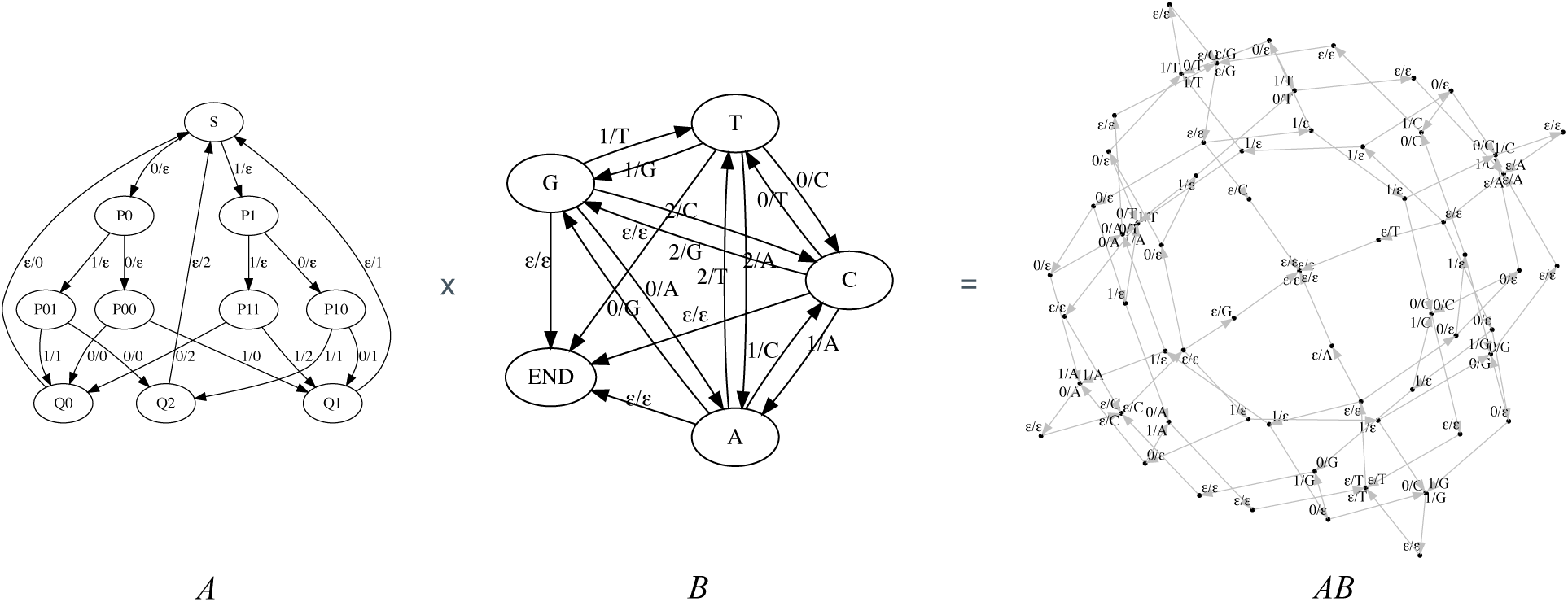
A nonrepeating DNA storage code can be factored into two separate state machines: one for converting binary sequences to ternary, and one for converting ternary to DNA [15]. In this diagram, a state machine transition is annotated *x/y* if it inputs *x* and outputs *y*; the symbol *ϵ* denotes the empty string. (*A*) Machine for (imperfectly) converting a binary input sequence into ternary, batching the input into groups of three binary digits and outputting pairs of ternary digits. This machine is inefficient in that the output is log(9)/ log(8) ≃1.06 times longer than it would be for a perfect conversion from base 2 to base 3 (because no triplet of input bits is ever converted the pair of output trits “22”, which means one of the nine possible output-trit pairs is wasted; more fundamentally, perfect conversion between indivisible integer bases is not possible with a finite state machine). In applications where the length of the input is not known in advance and so must be signaled with an end-of-file character (EOF), the ternary sequence “22” can be used to encode this EOF. (*B*) Machine for converting a ternary input sequence into a nonrepeating DNA sequence. The output of this machine is log(4)/ log(3) ≃1.26 times longer than the DNA would be if repeated nucleotides are allowed. (*AB*) Machine for a binary input sequence into a nonrepeating DNA sequence, obtained by “multiplying” *A* and *B*. The output of this machine is 4/3 times longer than the Shannon limit (obtained by multiplying the inefficiencies of the two constituent machines). The machine diagrams in this figure were generated automatically using Machine Boss and GraphViz.

**Fig. 2.**
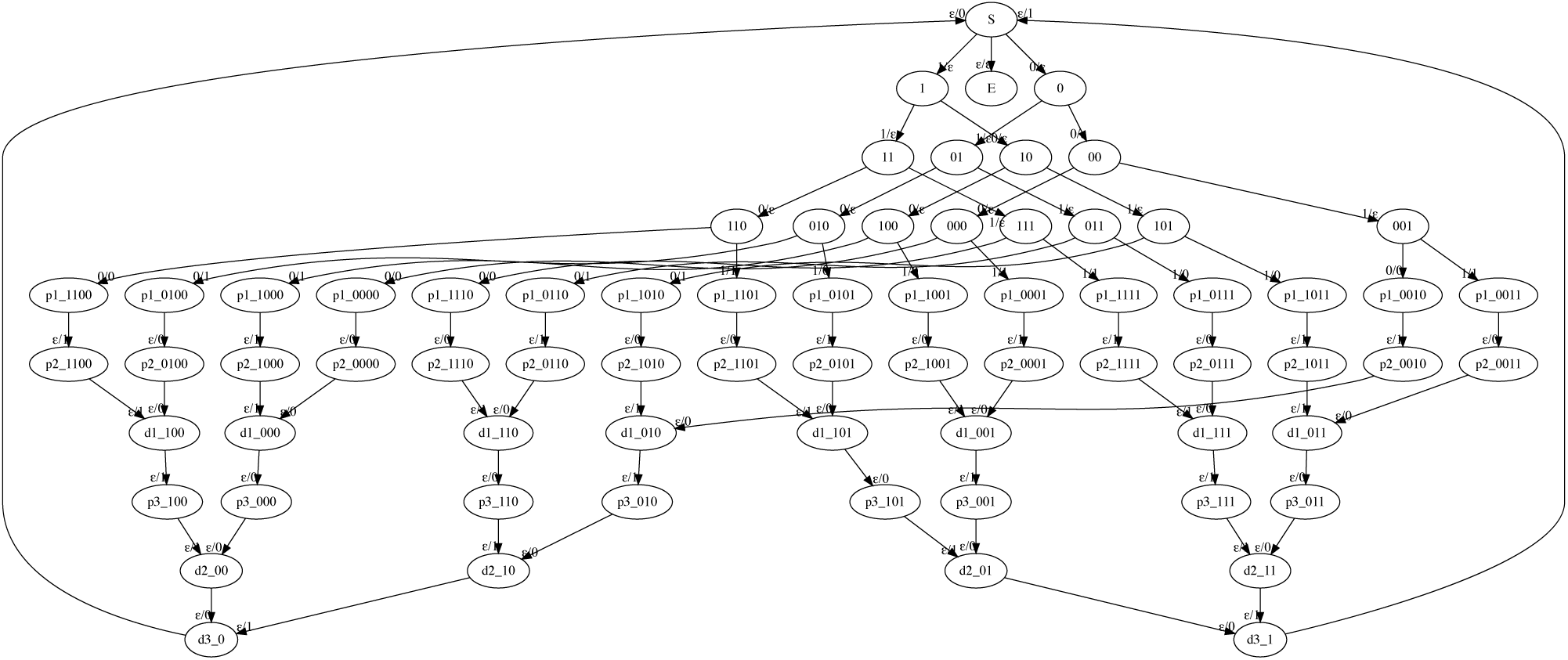
A state machine that implements the Hamming(7,4) error correction code, which interleaves 4 data bits with 3 parity bits, has 70 states and 85 transitions. A transition is annotated *x/y* if it inputs *x* and outputs *y*; the symbol *ϵ* denotes the empty string. The diagram was generated automatically using Machine Boss and GraphViz.

The approach of formally composing automata is well-documented in other applications of automata in computer science, for example in linguistics [33]. The automata, and the operations to combine or transform them, can be expressed compactly using the notation of linear algebra; in this view, an automaton formally represents an infinite matrix whose rows and columns are indexed by input and output sequences [5]. To take one example, the GeneWise combination of three translation, splicing, and error sub-machines corresponds straightforwardly to a three-way matrix multiplication. For some tasks, such as statistical phylogenetic alignment (where such automata generalize the idea of the “substitution matrix” to whole sequences, allowing indels as well as substitutions), this view of state machines as algebraic objects that can be systematically combined on the branches of a tree is absolutely central to the underlying probabilistic framework [20, 37, 45, 38, 49, 21]. Even without adopting the linear algebraic view, there is clear utility to being able to transform automata by simple operations like reverse-complementation or concatenation. Yet, for all the general-purpose state-machine libraries, this ability to formally operate on the state machines themselves is not generally available. Certainly, most libraries allow state machines to be constructed programmatically, by building appropriate data structures directly in the source code that links to the library. However, this is an intricate and error-prone procedure, and is a far cry from being able to construct state machines from modular components using reliable, general-purpose implementations of elementary operations such as “multiply”, “concatenate”, or “reverse-complement”.

Motivated by this gap in the bioinformatics tool chain, and finding ourselves repeatedly in need of reference implementations of automata-theoretic algorithms (for prototyping and debugging purposes in both HMM- and RNN-based applications) that allowed for algebraic manipulation of the underlying state machines, we developed Machine Boss, an open source software package that meets this need. In the Methods section and the Supplementary Information, we review the representation of state machines used throughout Machine Boss, and outline its capabilities. In the Results section, we describe nontrivial example applications of Machine Boss to several problems of interest; these include incorporating context-dependent error models (appropriate for nanopore sequencing) into GeneWise-like protein-to-DNA aligners, decoding the output of neural network basecallers for ONT sequencing instruments, and prototyping modular codes for encoding binary information in DNA. In the Discussion, we discuss how this sort of prototyping fits into a bioinformatics tool development workflow, and briefly mention several further applications.

## 2 RESULTS

### 2.1 Aligning protein sequences to nanopore reads with a context-dependent error model

Our first test of Machine Boss was an experiment to see whether richer error models might benefit a GeneWise-like protein-to-DNA search. Specifically, we sought to prototype an application to search amino acid sequences against individual nanopore reads, to see if they contained coding genes for known *E.coli* proteins. To do this, we combined a protein-to-DNA alignment model with two alternate error models: a “symmetric context-independent” error model parameterized by a substitution matrix and gap opening/extension parameters, and a richer “asymmetric context-dependent” error model with separate parameters for insertion and deletion (hence “asymmetric”), that also allows the error probabilities at a particular position of the genome to depend on the neighboring bases (hence “context-dependent”). For this application we were primarily interested in the power of the DNA substitution model; to reduce computation time, we did not incorporate an amino acid substitution model (as the analogous GeneWise algorithm does), but this can easily be incorporated.

The datasets and the construction and parameterization of the constituent state machines are described in more detail in the Methods and Supplementary Information. The protein-to-DNA model with symmetric context-independent errors has 242 states and 803 transitions (329 IO-conditioned). The model with asymmetric context-dependent errors has 1,349 states and 7,464 transitions (2,558 IO-conditioned).

After constructing the state machines, we used Machine Boss to generate custom C++ code for the Forward algorithm, compiled this code, and ran it to scan for a representative *E.coli* IS26 transposase protein (insB1, 167aa).

Figure 3 shows the results of this experiment. Broadly, both error models performed similarly, scoring on average around 1.4 bits per codon aligned. (This is a rather low alignment score, reflecting the extremely noisy nature of the training alignments.) We observed a small (3%) but significant improvement in log-odds scores for true positives when using the asymmetric context-dependent model, with negligible effect on log-odds scores for true negatives.

**Fig. 3.**
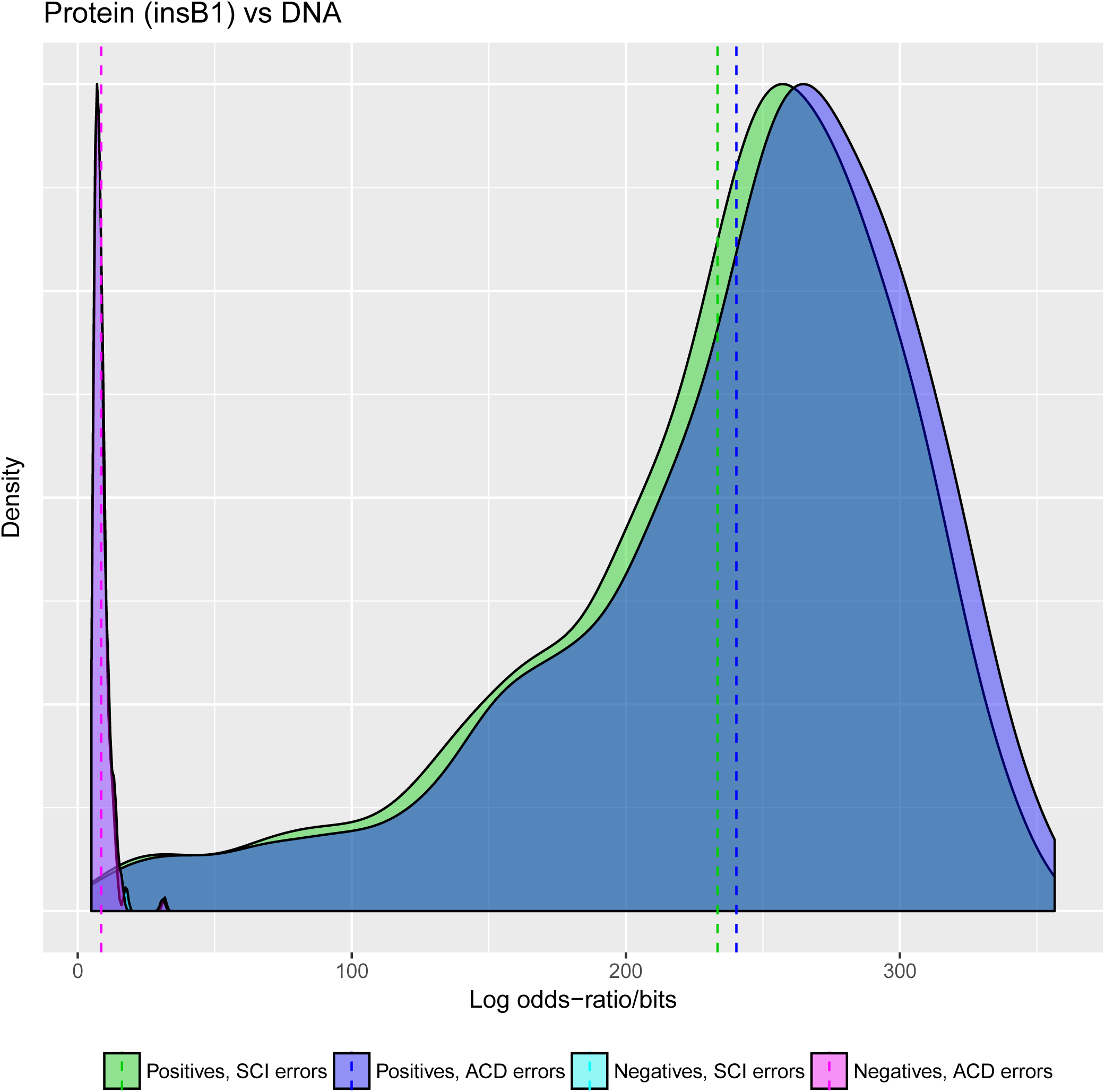
A richer error model slightly improves the discriminative power of a protein alignment to noisy sequencing reads. The plot shows the smoothed density of log-odds ratios of global alignments of *E.coli* protein insB1 to nanopore reads that fully contain a gene for that protein (“Positives”), versus those that don’t contain that protein or close homologs (“Negatives”), using error models with and without insertion/deletion asymmetry and context dependence (SCI = symmetric context-independent; ACD = asymmetric context-dependent). The log-odds ratio for a read is 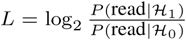 where ℋ_1_ is the hypothesis that the read contains the insB1 gene and ℋ_0_ the hypothesis that it does not. The mean of *L* is shown for each group. Using the asymmetric context-dependent error model increases the mean of the true positives (Δ*L* ≃ 6.9 bits, a relative increase of around 3%) with negligible effect on the true negatives (Δ*L* ≃ 0.1 bits).

To investigate how much of this improvement arose from the separate insertion and deletion probabilities, we prototyped a third error model, based on the symmetric context-independent model (and having similar state and transition counts), but relaxing the symmetry constraint between insertions and deletions. Results for this model are not shown in Figure 3 but its log-likelihoods for protein-to-DNA alignment generally lie in between the other two models, with a relative improvement of around 1% over the symmetric context-independent model. Thus, the improvement from allowing context-dependence appears to be roughly double the improvement from allowing symmetry-breaking, for this task.

### 2.2 Aligning protein domain profiles to nanopore reads

In the previous section we developed a state machine to model nanopore-specific sequencing errors and used that to better align protein sequences to nanopore reads. Machine Boss is able to combine arbitrary state machines, so the previous error models can be easily combined with other bioinformatics state machines. We next sought to investigate whether a richer error model would also benefit a profile HMM search. Instead of aligning a single protein sequence to a nanopore read, we combined a profile HMM with our nanopore specific error model and aligned it to the same set of nanopore reads from the previous section.

For this experiment we again focused on the insB1 transposase from the previous example and used the Pfam DDE_Tnp_IS1 domain (accession PF03400), which profiles its catalytic domain. We imported the HMMER-formatted profile HMM from the Pfam database into Machine Boss, and composed it with each of two error models: the symmetric context-independent error model, and the asymmetric context-dependent model. We did not use the asymmetric context-independent model in this experiment.

Results are shown in Figure 4. Proceeding from the symmetric context-independent model to the richer asymmetric context-dependent model, we see a relative increase in the log-likelihood of 4.1% for true positives and 2.2% for true negatives.

**Fig. 4.**
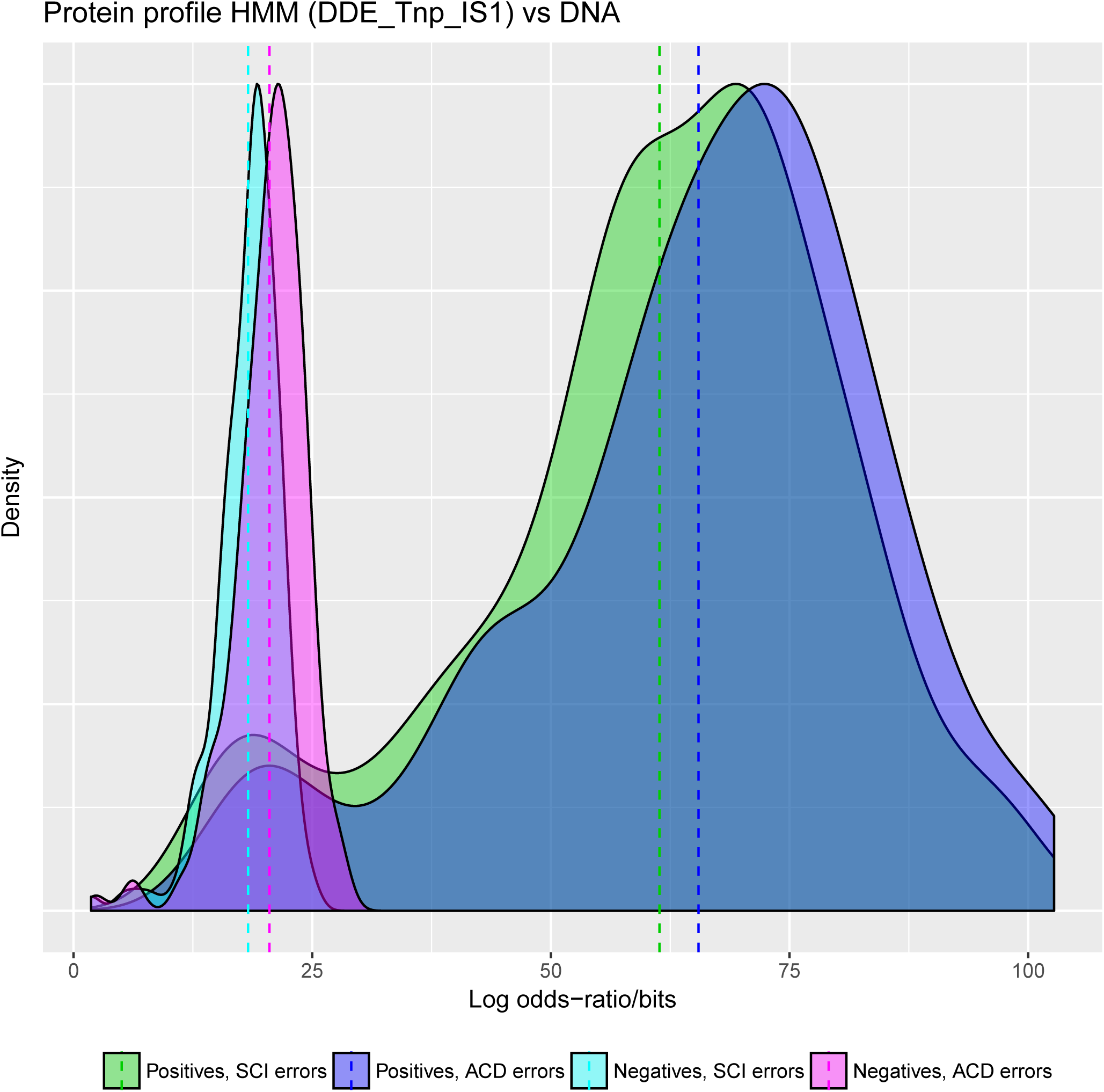
A richer error model slightly improves the discriminative power of a protein profile HMM search of noisy sequencing reads. The plot shows the smoothed density of log-odds ratios of the PFAM domain DDE_Tnp_IS1 (PF03400) for the IS1 transposase, aligned to nanopore ONT reads that fully contain a gene for the corresponding insB1 protein (“Positives”), versus those that don’t contain that protein or close homologs (“Negatives”), using error models with and without insertion/deletion asymmetry and context dependence (SCI = symmetric context-independent; ACD = asymmetric context-dependent). The log-odds ratio for a read is 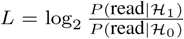 where ℋ_1_ is the hypothesis that the read contains the insB1 gene and ℋ_0_ the hypothesis that it does not. The mean of *L* is shown for each group. Using the asymmetric context-dependent error model increases the mean of the true positives (Δ*L* ≃ 4.1 bits) with a smaller effect on the true negatives (Δ*L* ≃ 2.2 bits).

#### 2.2.1 A note on code generation and performance

For this experiment we did not generate custom C++ code, for the reason that the Profile HMM-derived models are immense: even a relatively small model such as DDE_Tnp_IS1, which has 131 match states, grows to 50,766 states and 174,118 transitions (93,631 IO-conditioned) when combined with a DNA-to-protein model and the simplest nontrivial error model (symmetric context-independent). This in turn generates multiple C++ source files that are over 4Mb in size, including functions with over 50,000 lines of code (since each state requires at least one line), which in practice cause severe problems for C compilers (we were not able to compile these files using Apple LLVM version 9.0.0 / clang-900.0.39.2). It should yet be possible to compile these files by breaking up the generated code into smaller subroutines, at some small cost to performance; alternatively, it would be straightforward to generate efficient platform-specific assembly language directly, bypassing the C compiler altogether. We did not pursue either of these ideas, but instead for the profile HMM examples we used Machine Boss’s internal implementation of the Forward algorithm, which dynamically interprets the state machine (and so is generically re-usable for all models).

We would expect generic interpreted code to run slower than custom-generated compiled code. Empirically, we observe the compiled code runs around 40-fold faster than the interpreted code (e.g. on a 3.5GHz Intel Xeon E5, the compiled Forward algorithm runs at around 180M transitions/sec while the interpreted Forward algorithm runs at around 4M transitions/sec; these rates refer to the IO-conditioned transition counts).

### 2.3 Decoding the most likely output sequence of a neural network basecaller

Our third experiment tests the decoding algorithms used for basecalling on the Oxford Nanopore Technologies (ONT) sequencing platform. As a single strand of DNA (or RNA) passes through the protein nanopore, it perturbs an electrical current signal in a sequence-dependent way. A neural network trained with Connectionist Temporal Classification (CTC, [17]) outputs a probability distribution over sequences, which requires an additional decoding step to find the most likely sequence. CTC was developed for speech recognition and first applied to nanopore sequencing by Chiron [46], and was later adopted by various ONT basecallers.

Bonito (https://github.com/nanoporetech/bonito) is ONT’s most recent research basecaller: it uses a convolutional architecture based on QuartzNet [26], and is trained with CTC loss. In practice, Bonito uses Viterbi decoding, which simply takes the argmax of the logits and concatenates the resulting nucleotide and gap characters.

In this test, we compare the use of a Viterbi decoding scheme, which just finds the single most probable path through the data, with a beam search, a heuristic search algorithm which looks for the best label sequence. The CTC probability outputs are similar to profile HMMs and as such can be interpreted as state machines [42]. We evaluated these algorithms on a small sample of 100 reads from a publically available R9.4 *Klebsiella pneumoniae* dataset [50]. Accuracy was evaluated by aligning basecalled reads with minimap2 [28] to a reference genome from the same study. Each read was basecalled with a version of Bonito modified to save the network output, which was then loaded as a state machine into Machine Boss using the `--recognize-merge-csv` option, which constructs a state machine that merges repeated characters in the same manner as the CTC loss, as described in [17].

Results are shown in Figure 5. We found that the beam search yielded an increase of 0.2% median accuracy over Viterbi, though in practice its greater computational cost would likely not be worth such a slight improvement. These results were obtained with a beam width of 5; a larger beam size of 50 did not noticeably improve the results.

**Fig. 5.**
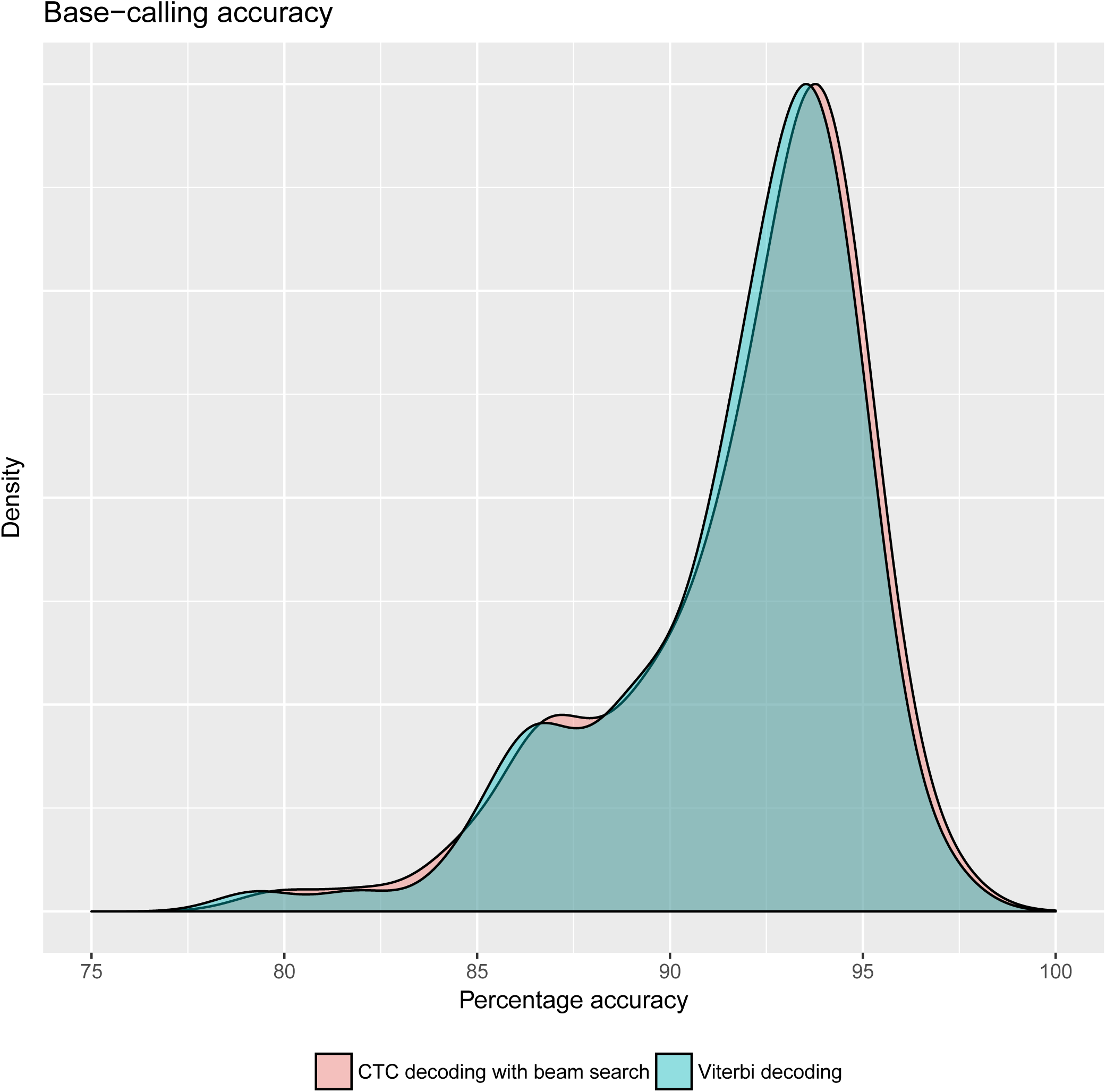
A beam-search decoding of the maximum likelihood sequence of Oxford Nanopore’s Bonito basecaller slightly outperforms a Viterbi best-path decoding on a sample of 100 *Klebsiella pneumoniae* reads. The percent accuracy is defined as the number of identities in the alignment divided by the total alignment length. Median accuracy with Viterbi was 92.8% while beam search yielded a median accuracy of 93.0%. This slight increase in accuracy does incur a computational cost: the beam search (width of 5) takes roughly 1.25 times as long as the Viterbi decoding. We further observe that a bespoke Python implementation of Viterbi decoding (optimized for this model architecture) was roughly 5 times as fast as Machine Boss’s generic C++ implementation of Viterbi decoding (which spends most of its time constructing and topo-sorting the state machine). This reinforces the conclusion that Machine Boss is better suited to prototyping and debugging during development, than to computationally intensive end-user applications.

In addition to the decoding of single reads, more elaborate dynamic programming algorithms for consensus decoding [42] can also easily be implemented and tested in Machine Boss. In this case performance is too slow for practical application to large datasets, though these reference implementations can be used to debug domain-specific software. In our case, Machine Boss has helped with testing our own consensus basecalling software PoreOver (https://github.com/jordisr/poreover, manuscript in preparation).

### 2.4 Constructing a repeat-avoiding code for DNA data storage

Our last computational experiment is motivated not by sequence analysis, but by analysis of the state machines themselves. We sought to investigate the complexity of error-tolerant codes for storing information in DNA.

We developed a state machine for converting binary information to DNA sequences using, as a starting point, the DNA storage code developed by Goldman *et al* [15]. The central idea of this code is that DNA homopolymers are often misread by many sequencing technologies, so this class of errors can be avoided altogether by never repeating a base in the encoded sequence. This leaves three nucleotides available for encoding information at any given position in the sequence, which corresponds to a radix-3 (ternary) representation.

We implement this as a multiplication of two machines: one that converts binary to ternary (with some unavoidable inflation of the message size), and a second machine that converts ternary to non-repeating DNA. This configuration is shown in Figure 1.

Using Machine Boss, we were able to successfully prototype this machine and to confirm that it accurately encodes binary messages to nonrepeating DNA strings and decodes in the opposite direction, with the ratio of binary to DNA message lengths asymptotically approaching the expected limit of 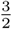. This ratio is calculated as follows. The conversion of binary to ternary is approximate, batching the input bits into triplets (with 2^3^ = 8 possibilities per batch) and outputting trits—i.e. ternary digits—in pairs (with 3^2^ = 9 possibilities per batch). Thus, the input sequence has 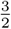 as many characters as the output. The ternary-to-DNA conversion converts each ternary digit to a single nucleotide, so the overall input/output ratio for the full binary-to-DNA conversion is also 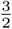.

This is slightly wasteful given that the Shannon information content of a nonrepeating DNA sequence is log_2_ 3 ≃ 1.58 bits/symbol, slightly greater than 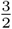. The wastage is incurred by the batched binary-to-ternary conversion, since there are more output possibilities than input possibilities for each batch. As can be seen in machine *A* of Figure 1, there is no triplet of input bits that will ever output the pair of trits “`22`”. This reflects a more general phenomenon that a finite-state machine cannot perfectly convert a radix-2 input to a radix-3 output (essentially, it can only compute the last digit of this conversion, which amounts to dividing the input by 3 and outputting the remainder; the quotient must then be fed into a similar machine to compute the second-last digit, and so on). It is possible to get quite close to the limit, though, by batching input bits in this way, and a batch size of 3 bits is a reasonably efficient compromise in terms of the number of states required by the machine; no improvement can be gained by improving the batch size until one reaches 11 bits, whereupon the ratio of input/output message lengths is 11/7 ≃ 1.57, but this requires 𝒪(2^11^) states to track each batch. Such a machine can readily be prototyped with Machine Boss, but the algorithms to manipulate and use the state machines become quite cumbersome for large machines (in addition to the well-known time complexity of dynamic programming to state machines, Machine Boss performs operations like topological sort and state elimination that can be slow for very large machines).

The input/output ratio of 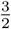 is approached asymptotically from below, because there is a necessary overhead involved in encoding the message length itself; our machine encodes this by employing the otherwise-unused pair of output trits “`22`” as an end-of-message terminator sequence. For simplicity, this mechanism is not included in Figure 1; when it is included, the combined machine for binary-to-nonrepeating-DNA conversion has 85 states and 132 transitions (44 IO-conditioned). The component machines were constructed with a short JavaScript program, and are available as presets in Machine Boss.

We can readily extend the above-described approach to study more elaborate DNA storage codes. For example, we can develop a DNA-encoding machine that avoids not just repeated nucleotides in the output, but also avoids certain nucleotide motifs, such as restriction enzyme sites; briefly, the transition graph of such a machine can be found by starting with a de Bruijn graph over *k*-mers, from which the prohibited *k*-mers are then deleted. (Of course, restriction enzyme sites that contain repeated nucleotides would already be excluded.) We might also incorporate error-correction units, such as Hamming codes or indel-resistant “watermarks”. Finally, we can incorporate technology-specific models of sequencing error, such as the nanopore error models described in previous sections, when decoding messages. All these variations can be implemented as modular machines and factored into the “matrix multiplication” of Figure 1. For example, introducing a Hamming(7,4) error-correcting parity code (Figure 2) to the nonrepeating-DNA code (Figure 1) yields a machine with 1,365 states and 1,812 transitions (292 IO-conditioned), whose input/output ratio is 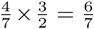. A deeper exploration of these ideas, using state machines to prototype the codes and investigating their error-correcting properties by simulation, is available in a separate preprint [19].

## 3 METHODS

Detailed descriptions of state machines and sequence data may be found in the Supplementary Information to this paper.

### 3.1 Weighted finite-state machines

The following definitions mostly parallel those found elsewhere [33, 49].

For our purposes, a *machine* is a tuple *T* = (Ω_*I*_, Ω_*O*_, Φ, *τ*, Θ, Ψ, *ν*) where Ω_*I*_ is an input alphabet, Ω_*O*_ is an output alphabet, Φ is a nonempty ordered list of states (of which the first element is the start state and the last is the end state), *τ* ⊆ Φ × (Ω_*I*_ ∪ {*ϵ*}) (Ω_*O*_ ∪ {*ϵ*}) × Φ is a set of transitions between states (labeled with input and/or output symbols), Θ is a set of named parameters (a subset of which are assigned nonnegative real values), Ψ is a set of constraints (partitioning Θ into rates, mutually exclusive probability, and other parameters), and *ν* : *τ* → Λ is the transition weight function, where Λ represents the set of closed-form differentiable expressions over Θ (with an expression grammar that allows real numbers, arithmetic operators, powers, exponentials and logarithms). Let ℳ denote the set of all such possible machines.

For a given input sequence 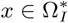 and output sequence 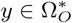, let *T*_*x,y*_ be the total weight of all paths through the transition graph of *T* that have input label *x* and output label *y*. The sequence weight *T*_*x,y*_ can be computed by the Forward algorithm in time 𝒪(|*x*| × |*y*|) and memory 𝒪(min(|*x*|, |*y*|)). The derivatives 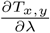 for *λ* ∈ Θ can be computed using the Forward-Backward algorithm. (Machine Boss implements these algorithm only for the case when all transition weights can be computed as real values, i.e. all relevant parameters are specified.)

This notation encourages us to think of *T* as being like an infinite matrix, indexed by sequences, with *T*_*x,y*_ being the element in row *x* and column *y*. We can then, for example, multiply machines like matrices: if *T, U* ∈ ℳ, then we can readily find a machine *TU* ∈ ℳ such that (*TU*)_*x,z*_ = ∑_*y*_ *T*_*x,y*_ *U*_*y,z*_. Other matrix expressions such as *T* + *U*, transpose(*T*), or *αT* (for some scalar *α*) are also straightforward to implement.

A machine for which Ω_*I*_ =ø is called a generator. A machine for which Ω_*O*_ =ø is called a recognizer. We can, for example, think of profile HMMs as generators (because they generate sequences as output) and regular expressions as recognizers (because they accept sequences as input). The labeling of one sequence as “input” and the other as “output” is arbitrary and, for many purposes, largely irrelevant. In the linear algebra analogy, the distinction between a generator and a recognizer corresponds to the choice as to whether to represent a vector in row or column form, and exchanging the “input” and “output” labels corresponds to taking the transpose.

### 3.2 Machine Boss capabilities

Machine Boss defines a (validatable) JSON format for machines in ℳ, and implements the following operations:

- Matrix-like operations such as multiplication, transposition, addition, intersection (a.k.a. point product or Hadamard product), the matrix identity (for a given alphabet), and multiplication by a scalar;
- String-like operations such as concatenation, reversal, reverse-complement, repetition, Kleene closure, local matching (padding with flanking states),
- HMM transition graph-related operations such as topological sorting, elimination of *ϵ*-transitions, elimination of redundant or inaccessible states, downsampling, normalization, and various probabilistic weightings;
- Construction of generators and recognizers for particular sequences, elementary patterns (e.g. wildcards), or regular expressions;
- Import of models from various sources such as HMMER files [13], FASTA files, CSV files, or HTTP fetches from PFAM [14] or DFAM [22];
- Export of models to GraphViz format;
- Useful built-in “preset” machines such as probabilistic Smith-Waterman [8], GeneWise-like models [3], DNA storage codes [15], the Jukes-Cantor model [25], and the Thorne-Kishino-Felstenstein model [47];
- Implementations of common dynamic programmic (Forward, Forward-Backward, Viterbi) and HMM-related algorithms (Baum-Welch and other forms of Expectation Maximization, respecting the user-specified parameter constraints), with banding heuristics;
- Implementation of search algorithms for finding the highest-weighted input or output sequences, or sampling such sequences probabilistically by weight; including prefix search, beam search, stochastic prefix search, MCMC, and simulated annealing;
- Generation of C++ code (32- or 64-bit) and/or JavaScript code for the Forward algorithm;
- Multiple convenient ways to specify input and output sequences (as strings from the command line, JSON arrays, or FASTA files);
- A flexible logging system for progress reports and debugging messages.

To compute the sum over all state paths, the Forward algorithm requires that the “silent” (i.e. *ϵ*-labeled) subset of the transition graph is acyclic and topologically sorted [12]. Most Machine Boss operations attempt to maintain this property in the state machines that they construct, automatically topo-sorting and eliminating silent cycles by marginalization. However, for large state machines (particularly when the transition weights are expressed symbolically as closed-form algebraic formulae, rather than as real numbers) these operations can become computationally expensive. For such cases, Machine Boss also offers inexact versions of the operations that either attempt to break silent cycles (by deleting silent transitions *i* → *j* where *j* < *i*, until no cycles remain) or just leave them in place (acknowledging that the Forward algorithm may then give a technically incorrect, albeit stable, result).

With the operations described, prototyping and evaluating new machines with Machine Boss is a relatively quick process that can take place interactively on the command line. In fact, many of these operations can be accessed in multiple ways: from the command line, via the JSON API, or by interfacing directly to the C++ API.

Machine Boss compiles on Apple Mac and Linux systems to a command-line executable with limited dependencies (GSL, and SSL if using the network capabilities) and can also be compiled to WebAssembly using emscripten (and thus run in the web browser, or in node).

## 4 DISCUSSION

Machine Boss can be useful for prototyping, testing, and theoretical analysis of state machines. In most cases, it is not suitable for developing polished bioinformatics tools, since further heuristic or custom optimizations of the generated state machines and code (beyond Machine Boss’s automated capabilities) is often possible and desirable.

As an example of this further optimization, our context-dependent error model has 50 states: a start state, an end state, and 48 states which consist of match, insert, and delete states repeated in 16 different flanking contexts. However, it is unnecessary to allocate storage for all 50 states during dynamic programming: the flanking context is always exactly determined by the position in the input genomic sequence, so only 5 states are ever accessible at any position in the dynamic programming matrix. An optimized implementation could make use of this, but Machine Boss currently lacks the sophistication to deduce such optimizations automatically. Rather, Machine Boss can be used (as we have done here) to evaluate whether such development is worthwhile, and to provide a robust reference implementation against which the results of a more optimized version can be checked.

In this manner, we have also found Machine Boss useful for debugging deep learning algorithms, such as beam-search decoding of RNNs [42] The outputs of ONT’s recent RNN basecallers can be interpreted as transition weights in a finite-state machine [24, 50]. In analyzing and improving on these results, we have found Machine Boss useful as a debugging and profiling tool.

Compared to recent deep learning approaches, automata retain some merits: they are highly interpretable, conceptually straightforward, and generally predictable. The interpretability is especially appealing when paths through the automaton have clear meaning—as is the case when state machines are used to represent biological processes such as translation and splicing, information-theoretic processes like radix-based coding, or evolutionary processes such as indels (for which purpose Machine Boss includes a reference implementation of the Thorne-Kishino-Felsenstein model [47]). The software development was motivated directly by these cases, but the algorithms implemented are general enough that we have been able to use it for applications in nanopore analysis as well. The README file in the Machine Boss repository describes several further applications, including machines to search for a PROSITE regular expression in a protein sequence and to count copies of this motif in a (translated) DNA sequence. As with the examples in this paper, the power of this approach rests on the ability to combine such state machines in a general way, together with new machines as yet undeveloped.

## Supporting information

Models

## 5 ACKNOWLEDGMENTS

We thank Ewan Birney and Tim Massingham for informative discussions. This work was supported by NIH/NCI grant U24-CA220441, NIH/NHGRI grant R01-HG004483, NIH/NIGMS grant R01-GM080203, and by a research gift from Oxford Nanopore Technologies.

## REFERENCES

[1] M. Abadi, A. Agarwal, P. Barham, E. Brevdo, Z. Chen, C. Citro, Greg S. Corrado, A. Davis, J. Dean, M. Devin, S. Ghemawat, I. Goodfellow, A. Harp, G. Irving, M. Isard, Y. Jia, R. Jozefowicz, L. Kaiser, M. Kudlur, J. Levenberg, D. Mané, R. Monga, S. Moore, D. Murray, C. Olah, M. Schuster, J. Shlens, B. Steiner, I. Sutskever, K. Talwar, P. Tucker, V. Vanhoucke, V. Vasudevan, F. Viégas, O. Vinyals, P. Warden, M. Wattenberg, M. Wicke, Y. Yu, and X. Zheng. TensorFlow: Large-scale machine learning on heterogeneous systems, 2015. Software available from tensorflow.org.

[2] M. Alexandersson, S. Cawley, and L. Pachter. SLAM: cross-species gene finding and alignment with a generalized pair hidden Markov model. Genome Research, 13(3):496–502, 2003.

[3] E. Birney, M. Clamp, and R. Durbin. GeneWise and GenomeWise. Genome Research, 14(5):988–995, 2004.

[4] E. Birney and R. Durbin. Dynamite: a flexible code generating language for dynamic programming methods used in sequence comparison. In T. Gaasterland, P. Karp, K. Karplus, C. Ouzounis, C. Sander, and A. Valencia, editors, Proceedings of the Fifth, pages 56–64, Menlo Park, CA, 1997. AAAI Press.

[5] A. Bouchard-Côté. A note on probabilistic models over strings: the linear algebra approach. Bulletin of Mathematical Biology, 75(12):2529–2550, Dec 2013.

[6] V. Boža, B. Brejová, and T. Vinař. DeepNano: Deep recurrent neural networks for base calling in MinION nanopore reads. PLoS ONE, 12(6):e0178751, 2017.

[7] M. Brown, R. Hughey, A. Krogh, I. S. Mian, K. Sjölander, and D. Haussler. Using Dirichlet mixture priors to derive hidden Markov models for protein families. In L. Hunter, D. B. Searls, and J. Shavlik, editors, Proceedings of the First, pages 47–55, Menlo Park, CA, 1993. AAAI Press.

[8] P. Bucher and K. Hofmann. A sequence similarity search algorithm based on a probabilistic interpretation of an alignment scoring system. In D. J. States, P. Agarwal, T. Gaasterland, L. Hunter, and R. F. Smith, editors, Proceedings of the Fourth, pages 44–51, Menlo Park, CA, 1996. AAAI Press.

[9] C. Burge and S. Karlin. Prediction of complete gene structures in human genomic DNA. Journal of Molecular Biology, 268(1):78–94, 1997.

[10] M. David, L. J. Dursi, D. Yao, P. C. Boutros, and J. T. Simpson. Nanocall: an open source basecaller for Oxford Nanopore sequencing data. Bioinformatics, 33(1):49–55, Jan 2017.

[11] C. B. Do, M. S. P. Mahabhashyam, M. Brudno, and S. Batzoglou. ProbCons: Probabilistic consistency-based multiple sequence alignment. Genome Research, 15(2):330–340, Feb 2005. Comparative Study.

[12] R. Durbin, S. Eddy, A. Krogh, and G. Mitchison. Biological Sequence Analysis: Probabilistic Models of Proteins and Nucleic Acids. Cambridge University Press, Cambridge, UK, 1998.

[13] S. R. Eddy. A new generation of homology search tools based on probabilistic inference. Genome Inform, 23(1):205–211, Oct 2009.

[14] S. El-Gebali, J. Mistry, A. Bateman, S. R. Eddy, A. Luciani, S. C. Potter, M. Qureshi, L. J. Richardson, G. A. Salazar, A. Smart, E. L. L. Sonnhammer, L. Hirsh, L. Paladin, D. Piovesan, S. C. E. Tosatto, and R. D. Finn. The Pfam protein families database in 2019. Nucleic Acids Res., 47(D1):D427–D432, Jan 2019.

[15] N. Goldman, P. Bertone, S. Chen, C. Dessimoz, E. M. LeProust, B. Sipos, and E. Birney. Towards practical, high-capacity, low-maintenance information storage in synthesized DNA. Nature, 494(7435):77–80, 2013.

[16] O. Gotoh. An improved algorithm for matching biological sequences. Journal of Molecular Biology, 162:705–708, 1982.

[17] A. Graves, S. Fernández, F. Gomez, and J. Schmidhuber. Connectionist temporal classification: Labelling unsegmented sequence data with recurrent neural networks. In Proceedings of the 23rd International Conference on Machine Learning, ICML ‘06, pages 369–376, New York, NY, USA, 2006. ACM.

[18] M. Gribskov, A. D. McLachlan, and D. Eisenberg. Profile analysis: detection of distantly related proteins. Proceedings of the National Academy of Sciences of the USA, 84:4355–4358, 1987.

[19] I. Holmes. Modular non-repeating codes for DNA storage. CoRR, abs/1606.01799, 2016.

[20] I. Holmes and W. J. Bruno. Evolutionary HMMs: a Bayesian approach to multiple alignment. Bioinformatics, 17(9):803–820, 2001.

[21] I. H. Holmes. Historian: accurate reconstruction of ancestral sequences and evolutionary rates. Bioinformatics, 33(8):1227–1229, 04 2017.

[22] R. Hubley, R. D. Finn, J. Clements, S. R. Eddy, T. A. Jones, W. Bao, A. F. Smit, and T. J. Wheeler. The Dfam database of repetitive DNA families. Nucleic Acids Res., 44(D1):D81–89, Jan 2016.

[23] M. Jain, I. T. Fiddes, K. H. Miga, H. E. Olsen, B. Paten, and M. Akeson. Improved data analysis for the MinION nanopore sequencer. Nat. Methods, 12(4):351–356, Apr 2015.

[24] M. Jain, S. Koren, K. H. Miga, J. Quick, A. C. Rand, T. A. Sasani, J. R. Tyson, A. D. Beggs, A. T. Dilthey, I. T. Fiddes, S. Malla, H. Marriott, T. Nieto, J. O’Grady, H. E. Olsen, B. S. Pedersen, A. Rhie, H. Richardson, A. R. Quinlan, T. P. Snutch, L. Tee, B. Paten, A. M. Phillippy, J. T. Simpson, N. J. Loman, and M. Loose. Nanopore sequencing and assembly of a human genome with ultra-long reads. Nat. Biotechnol., 36(4):338–345, 04 2018.

[25] T. H. Jukes and C. Cantor. Evolution of protein molecules. In Mammalian Protein Metabolism, pages 21–132. Academic Press, New York, 1969.

[26] S. Kriman, S. Beliaev, B. Ginsburg, J. Huang, O. Kuchaiev, V. Lavrukhin, R. Leary, J. Li, and Y. Zhang. QuartzNet: Deep Automatic Speech Recognition with 1D Time-Channel Separable Convolutions. pages 2–6, 2019.

[27] T. Y. Lam and I. M. Meyer. HMMCONVERTER 1.0: a toolbox for hidden Markov models. Nucleic Acids Res., 37(21):e139, Nov 2009.

[28] H. Li. Minimap2: pairwise alignment for nucleotide sequences. Bioinformatics, 34(18):3094–3100, 09 2018.

[29] P. C. Lott and I. Korf. StochHMM: a flexible hidden Markov model tool and C++ library. Bioinformatics, 30(11):1625–1626, Jun 2014.

[30] A. Löytynoja and N. Goldman. An algorithm for progressive multiple alignment of sequences with insertions. Proceedings of the National Academy of Sciences of the USA, 102(30):10557–62, 2005.

[31] G. Lunter. HMMoC–a compiler for hidden Markov models. Bioinformatics, 23(18):2485–2487, Sep 2007.

[32] I. M. Meyer and R. Durbin. Gene structure conservation aids similarity based gene prediction. Nucleic Acids Research, 32(2):776–783, 2004.

[33] M. Mohri, F. Pereira, and M. Riley. Weighted finite-state transducers in speech recognition. Computer Speech and Language, 16(1):69–88, 2002.

[34] E. F. Moore. Gedanken-experiments on sequential machines. In C. Shannon and J. McCarthy, editors, Automata Studies, pages 129–153. Princeton University Press, Princeton, NJ, 1956.

[35] S. B. Needleman and C. D. Wunsch. A general method applicable to the search for similarities in the amino acid sequence of two proteins. Journal of Molecular Biology, 48:443–453, 1970.

[36] D. K. Ralph and F. A. Matsen. Consistency of VDJ Rearrangement and Substitution Parameters Enables Accurate B Cell Receptor Sequence Annotation. PLoS Comput. Biol., 12(1):e1004409, Jan 2016.

[37] B. D. Redelings and M. A. Suchard. Joint Bayesian estimation of alignment and phylogeny. Systematic Biology, 54(3):401–418, 2005.

[38] B. D. Redelings and M. A. Suchard. Incorporating indel information into phylogeny estimation for rapidly emerging pathogens. BMC Evolutionary Biology, 7:40, 2007.

[39] A. Schliep, W. Georgi, W. Rungsarityotin, I. G. Costa, and A. Schönhuth. The general hidden markov model library: Analyzing systems with unobservable states. Proceedings of the Heinz-Billing-Price, 01 2004.

[40] A. Siepel and D. Haussler. Phylogenetic estimation of context-dependent substitution rates by maximum likelihood. Molecular Biology and Evolution, 21(3):468–488, 2004.

[41] A. Siepel, K. S. Pollard, and D. Haussler. New methods for detecting lineage-specific selection. In Research in Computational Molecular Biology, pages 190–205, Berlin, Heidelberg, 2006. Springer Berlin Heidelberg.

[42] J. Silvestre-Ryan and I. Holmes. Consensus Decoding of Recurrent Neural Network Basecallers, pages 128–139. 01 2018.

[43] G. S. Slater and E. Birney. Automated generation of heuristics for biological sequence comparison. BMC Bioinformatics, 6:31, Feb 2005.

[44] T. F. Smith and M. S. Waterman. Identification of common molecular subsequences. Journal of Molecular Biology, 147:195–197, 1981.

[45] M. A Suchard and B. D Redelings. BAli-Phy: simultaneous Bayesian inference of alignment and phylogeny. Bioinformatics, 22(16):2047–2048, Aug 2006.

[46] H. Teng, M. B. Hall, T. Duarte, M. Duc Cao, and L. Coin. Chiron: Translating nanopore raw signal directly into nucleotide sequence using deep learning. bioRxiv, 2017.

[47] J. L. Thorne, H. Kishino, and J. Felsenstein. An evolutionary model for maximum likelihood alignment of DNA sequences. Journal of Molecular Evolution, 33:114–124, 1991.

[48] P. Veličković and P. Liò. Muxstep: an open-source C ++ multiplex HMM library for making inferences on multiple data types. Bioinformatics, 32(16):2562–2564, 08 2016.

[49] O. Westesson, G. Lunter, B. Paten, and I. Holmes. Accurate reconstruction of insertion-deletion histories by statistical phylogenetics. PLoS ONE, 7(4):e34572, 2012.

[50] R. R. Wick, L. M. Judd, and K. E. Holt. Performance of neural network basecalling tools for Oxford Nanopore sequencing. Genome Biol., 20(1):129, 06 2019.

