## Supplementary Information for "Machine Boss: Rapid Prototyping of Bioinformatic Automata"

### 1 State machines

In this section we briefly describe the machines developed for the experiments in this paper, which we also make available as supplementary files.

For each model, we describe the number of states ($S$) and transitions ($T$). These are directly relevant to estimating memory and time complexity. The memory complexity of dynamic programming to sequences of lengths $L, M$ is $\mathcal{O}(S \min L, M)$, or $\mathcal{O}(SLM)$ if traceback is required; checkpointing approaches can reduce the memory required for traceback [2], but are currently not implemented by Machine Boss. The time complexity is $\mathcal{O}(TLM)$. Machine Boss does some optimization to ensure that only the subset of transitions compatible with a given input and output character is considered at any given cell in the dynamic programming matrix; if we consider the maximum size of this subset (taken over all input/output character combinations), the effective number of transitions is $T_{\mathrm{IO}}$ and the time complexity is $\mathcal{O}(T_{\mathrm{IO}}LM)$. We refer to this latter transition count $T_{\mathrm{IO}}$ as the number of "IO-conditioned transitions". (In principle Machine Boss could optimize even further and consider IO-conditioned states, or conditioning that used more than one character of IO context; in practice we have not implemented such schemes.)

Note that the transition counts $T$ for Machine Boss are not directly comparable to those of GeneWise. One reason for discrepancies is that GeneWise (along with other standard bioinformatics automata [1]) adopts conventions that reduce the apparent number of transitions, including associating outputs with states instead of transitions and collapsing transitions with the same source and destination states (but different input or output labels). This difference is partially addressed by using IO-conditioned transition counts ($T_{\mathrm{IO}}$ instead of $T$). Another reason for discrepancies is that Machine Boss imposes a constraint that transition input and output labels can have at most one character, whereas GeneWise allows strings. This is a hard requirement of Machine Boss (necessary for simple manipulation of machines).

### 1.1 Symmetric context-independent read error model

The model that we refer to as "symmetric context-independent" is a conditionally-normalized pair HMM (i.e. a transducer) with a $4 \times 4$ substitution probability matrix, and gap opening and extension probabilities (identical for both insertions and deletions; hence "symmetric"). (This is somewhat simplified compared to the transducer fit to nanopore data by Jain *et al* [3] which allowed separate insertion and deletion probabilities, and further distinguished between short and long indels; that is, insertion and deletion lengths are each distributed according to a two-component mixture of geometric distributions.) This machine is available as a preset in Machine Boss; it has 8 states and 34 transitions (11 IO-conditioned).

Using Machine Boss, we trained this model on the 170-read training set using Expectation Maximization, conditioned on the read-to-reference alignments.

### 1.2 Asymmetric context-dependent read error model

The model we call "asymmetric context-dependent" is also a transducer, but allows separate probability distributions for insertions and deletions (though these are still single-component geometric distributions); this is the basis for calling it "asymmetric". Further, all substitution and indel probabilities in this model are allowed to depend on the two flanking nucleotides in the ancestral sequence; this is its "context-dependence". The machine was generated using a short JavaScript program; it has 50 states and 1,152 transitions (112 IO-conditioned).

We trained this model on the same 170-read, 1.7M-base training set and evaluated it on the same 165-read, 1.5M-base test set.

Under the asymmetric context-dependent error model, the log-likelihood of the training set increases by 1.14% and the test set log-likelihood increases by 0.84%, relative to the analogous log-likelihoods for the symmetric context-independent model.

### 1.3 Asymmetric context-independent read error model

To investigate how much of the observed likelihood improvement was due to allowing context-dependent parameters and how much was due to allowing separate insertion and deletion probabilities, we prototyped a third read error model, the "asymmetric context-independent" model, essentially identical to the "symmetric context-independent" model (in terms of states and transitions) but allowing the insertion and deletion opening and extension probabilities to be fitted independently.

Compared to the symmetric context-independent model, this model showed a relative log-likelihood improvement of 0.5% on the training set and 0.4% on the test set. Together with the other results, this suggests that the improvement from allowing context-dependence of error parameters is real, and slightly greater than the improvement arising from symmetry-breaking between insertions and deletions.

#### 1.4 Protein-to-DNA alignment

Our protein-to-DNA models were obtained by "multiplying" a reverse-translation model (which accepts a protein sequence as input, and emits a DNA sequence as output, using the *E.coli* genetic code and codon frequencies) with one of the two read error models, dividing nucleotide output weights by 1/4 to yield an odds-ratio compared to a uniform null model for DNA, and concatenating this with unit-weight flanking models to allow local alignment.

Constructing this machine is a one-line recipe in Machine Boss, given that a machine for reverse-translation using the standard genetic code is available as a built-in preset.

The protein-to-DNA machine has 84 states and 144 transitions (23 IO-conditioned).

#### 1.5 Profile-to-DNA alignment

Our profile-to-DNA models were obtained by "multiplying" a reverse-translation model (which accepts a protein sequence as input, and emits a DNA sequence as output, using the *E.coli* genetic code and codon frequencies) with one of the two read error models, dividing nucleotide output weights by 1/4 to yield an odds-ratio compared to a uniform null model for DNA, and concatenating this with unit-weight flanking models to allow local alignment.

As with the protein-to-DNA machine, constructing in Machine Boss is a one-line recipe, employing the built-in functionality to construct machines from HMMER profiles. In general, a HMMER model with length $L$ for an alphabet of size $A$ is converted to a Machine Boss model with $5L + 4$ states and $(2A + 7)L + A + 3$ transitions ($9L + 4$ IO-conditioned). Thus, the DDE_Tnp_IS1 profile that we used in this paper, which is 131 sub-units long, has 659 states and 6,180 transitions (1,183 IO-conditioned).

### 2 Sequence data

For the work in this paper we used the *E.coli* K-12 MG1655 genome sequence obtained by Loman *et al* using the ONT MinION sequencer with R7.3 nanopore chemistry and the Metrichor basecaller [6]. We aligned the raw basecalled reads to the *E.coli* genome using minimap2 [5]. The experimental setup (R7.3 chemistry, Metrichor basecaller) is reported to yield 78-85% accuracy; when we realigned reads using minimap2, we observed a sequence identity closer to 65%, perhaps due to errors in realignment. More recent nanopore chemistries and basecallers have a significantly lower error rate [7, 4, 8]. However, since our purpose was a bioinformatics exercise to demonstrate that automata can be used as error models for noisy sequence reads, we proceeded to use the 2015 data of Loman *et al* even though higher-accuracy nanopore sequence has since become available.

We selected a subsample of 170 read alignments as a training set, sampling the reads quasi-randomly (by taking one read in every thousand). This training set contained 1.7M total read bases, aligned to 1.8M genomic bases. We further selected 165 reads as a test set (disjoint from the training set) containing 1.5M total read bases aligned to 1.6M genomic bases.

The tests that involved searching for protein-coding gene homology, require a set of Negative and Positive controls. The "Positives" are reads known (from the *E.coli* genome annotation) to contain the insB1 gene, whereas the "Negatives" must not contain it. For the negatives, we started with the test set and excluded any reads that overlapped with insB1 (or any close homologs, e.g. other transposases).

### References


[1] R. Durbin, S. Eddy, A. Krogh, and G. Mitchison. *Biological Sequence Analysis: Probabilistic Models of Proteins and Nucleic Acids.* Cambridge University Press, Cambridge, UK, 1998.

[2] D. S. Hirschberg. A linear space algorithm for computing maximal common subsequences. *Communications of the ACM*, 18:341–343, 1975.

[3] M. Jain, I. T. Fiddes, K. H. Miga, H. E. Olsen, B. Paten, and M. Akeson. Improved data analysis for the MinION nanopore sequencer. *Nat. Methods*, 12(4):351–356, Apr 2015.

[4] M. Jain, S. Koren, K. H. Miga, J. Quick, A. C. Rand, T. A. Sasani, J. R. Tyson, A. D. Beggs, A. T. Dilthey, I. T. Fiddes, S. Malla, H. Marriott, T. Nieto, J. O'Grady, H. E. Olsen, B. S. Pedersen, A. Rhie, H. Richardson, A. R. Quinlan, T. P. Snutch, L. Tee, B. Paten, A. M. Phillippy, J. T. Simpson, N. J. Loman, and M. Loose. Nanopore sequencing and assembly of a human genome with ultra-long reads. *Nat. Biotechnol.*, 36(4):338–345, 04 2018.

[5] H. Li. Minimap2: pairwise alignment for nucleotide sequences. *Bioinformatics*, 34(18):3094–3100, 09 2018.

[6] N. J. Loman, J. Quick, and J. T. Simpson. A complete bacterial genome assembled de novo using only nanopore sequencing data. *Nat. Methods*, 12(8):733–735, Aug 2015.

[7] H. Teng, M. B. Hall, T. Duarte, M. Duc Cao, and L. Coin. Chiron: Translating nanopore raw signal directly into nucleotide sequence using deep learning. *bioRxiv*, 2017.

[8] R. R. Wick, L. M. Judd, and K. E. Holt. Performance of neural network basecalling tools for Oxford Nanopore sequencing. *Genome Biol.*, 20(1):129, 06 2019.
