## Supplementary material for "Machine Boss: Rapid Prototyping of Bioinformatic Automata": Models: asymmetric_context_dependent_error_model_generator.js

```
#!/usr/bin/env node

var fs = require('fs'),
    Getopt = require('node-getopt')

var getopt = Getopt.create([
    ['a' , 'alphabet=STRING'  , 'alphabet'],
    ['n' , 'name=STRING'      , 'name'],
    ['h' , 'help'             , 'display this help message']
])              // create Getopt instance
.bindHelp()     // bind option 'help' to default action
var opt = getopt.parseSystem() // parse command line

var alph = (opt.options.alphabet || "ACGT").split("")
var name = opt.options.name || "dna2"

function not (param) {
    return { "-": [true, param] }
}

function times (expr) {
  return Array.prototype.slice.call (arguments, 1).reduce (function (expr1, expr2) {
    return { "*": [expr1, expr2] }
  }, expr)
}

function sum (expr) {
  return Array.prototype.slice.call (arguments, 1).reduce (function (expr1, expr2) {
    return { "+": [expr1, expr2] }
  }, expr)
}

var startState = 'start', endState = 'end'
var startStateInfo = { id: startState, trans: [] },
    endStateInfo = { id: endState }

function matchState (leftContext, rightContext) { return 'mat' + leftContext + rightContext }
function insertState (leftContext, rightContext) { return 'ins' + leftContext + rightContext }
function deleteState (leftContext, rightContext) { return 'del' + leftContext + rightContext }

function eqmProb (i) { return 'eqm' + i }
function subProb (i, j, leftContext, rightContext) { return 'pSub' + i + j + '_' + leftContext + rightContext }
function insOpen (leftContext, rightContext) { return 'pInsOpen' + '_' + leftContext + rightContext }
function insExtend (leftContext, rightContext) { return 'pInsExt' + '_' + leftContext + rightContext }
function insChar (i, leftContext, rightContext) { return 'pInsChar' + i + '_' + leftContext + rightContext }
function delOpen (leftContext, rightContext) { return 'pDelOpen' + '_' + leftContext + rightContext }
// function delExtend (leftContext, rightContext) { return 'pDelExt' + '_' + leftContext + rightContext }
function delChar (j, leftContext, rightContext) { return 'pDelChar' + j + '_' + leftContext + rightContext }

function insOpenChar (i, leftContext, rightContext) { return times (insOpen (leftContext, rightContext), insChar (i, leftContext, rightContext)) }
function insExtendChar (i, leftContext, rightContext) { return times (insExtend (leftContext, rightContext), insChar (i, leftContext, rightContext)) }
function delOpenChar (j, leftContext, rightContext) { return times (delOpen (leftContext, rightContext), delChar (j, leftContext, rightContext)) }
// function delExtendChar (j, leftContext, rightContext) { return times (delExtend (leftContext, rightContext), delChar (j, leftContext, rightContext)) }
function delExtendChar (j, leftContext, rightContext) { return delChar (j, leftContext, rightContext) }

var states = [startStateInfo]
var norms = [], probs = []
alph.forEach (function (l) {
  alph.forEach (function (r) {
    // declare states & transitions
    startStateInfo.trans.push ({ to: matchState(l,r),
                                 weight: eqmProb(l) })
    states.push ({ id: matchState(l,r),
                   trans: (alph.reduce (function (list, c) {
                     return list.concat (alph.map (function (d) {
                       return { to: matchState(r,c),
                                in: r,
                                out: d,
                                weight: times (not(delOpenChar(r,l,c)), not(insOpen(l,r)), subProb(r,d,l,c)) }
                     })).concat ([ { to: deleteState(r,c),
                                     in: r,
                                     weight: delOpenChar(r,l,c) },
                                   { to: insertState(l,r),
                                     out: c,
                                     weight: times (not(delOpenChar(r,l,c)), insOpenChar(c,l,r)) } ])
                   }, [{ to: endState,
                         weight: eqmProb(r) }])) },
                 { id: insertState(l,r),
                   trans: (alph.reduce (function (list, c) {
                     return list.concat (alph.map (function (d) {
                       return { to: matchState(r,c),
                                in: r,
                                out: d,
                                weight: times (not(insExtend(l,r)), subProb(r,d,l,c)) }
                     })).concat ([ { to: insertState(l,r),
                                     out: c,
                                     weight: insExtendChar(c,l,r) } ])
                   }, [{ to: endState,
                         weight: times (not(insExtend(l,r)), eqmProb(r)) }])) },
                 { id: deleteState(l,r),
                   trans: (alph.reduce (function (list, c) {
                     return list.concat (alph.map (function (d) {
                       return { to: matchState(r,c),
                                in: r,
                                out: d,
                                weight: times (not(delExtendChar(r,l,c)), not(insOpen(l,r)), subProb(r,d,l,c)) }
                     })).concat ([ { to: deleteState(r,c),
                                     in: r,
                                     weight: delExtendChar(r,l,c) },
                                   { to: insertState(l,r),
                                     out: c,
                                     weight: times (not(delExtendChar(r,l,c)), insOpenChar(c,l,r)) } ])
                   }, [{ to: endState,
                         weight: eqmProb(r) }])) })
    // declare parameter groups
    alph.forEach (function (c) {
      norms.push (alph.map (function (d) { return subProb(c,d,l,r) }))
    })
    norms.push (alph.map (function (c) { return insChar(c,l,r) }))
    probs.push (insOpen(l,r), insExtend(l,r), delOpen(l,r) /* , delExtend(l,r) */ )
    probs = probs.concat (alph.map (function (c) { return delChar(c,l,r) }))
  })
})
states.push (endStateInfo)
norms.push (alph.map (function (c) { return eqmProb(c) }))

var machine = { state: states,
                cons: { norm: norms,
                        prob: probs } }

console.log (JSON.stringify (machine, null, 2))
```
